# TXNIP loss expands Myc-dependent transcriptional programs by increasing Myc genomic binding

**DOI:** 10.1101/2022.08.04.502753

**Authors:** Tian-Yeh Lim, Blake R. Wilde, Mallory L. Thomas, Kristin E. Murphy, Jeffery M. Vahrenkamp, Megan E. Conway, Katherine E. Varley, Jason Gertz, Donald E. Ayer

**Author notes:** Department of Biological Chemistry, David Geffen School of Medicine, University of California, Los Angeles, Los Angeles, CA 90095, USA. Department of Biomedical Genetics, Wilmot Cancer Institute, University of Rochester Medical Center, Rochester, NY, USA.

## Abstract

*c-Myc* protooncogene places a demand on glucose uptake to drive glucose-dependent biosynthetic pathways. To achieve this demand, c-Myc protein (Myc henceforth) drives the expression of glucose transporters and represses the expression of Thioredoxin Interacting Protein (TXNIP), which is a potent negative regulator of glucose uptake. A Myc_high_/TXNIP_low_ gene signature is clinically significant as it correlates with poor clinical prognosis in Triple-Negative Breast Cancer (TNBC) but not in other subtypes of breast cancer. To better understand how TXNIP function contributes to the aggressive behavior of TNBC, we generated TXNIP null MDA-MB-231 (231:TKO) cells for our study. We show here that TXNIP loss drives a transcriptional program that resembles those driven by Myc and increases global Myc genome occupancy. TXNIP loss allows Myc to invade the promoters and enhancers of target genes that are potentially relevant to cell transformation. Together, these findings suggest that TXNIP is a broad repressor of Myc genomic binding. The increase in Myc genomic binding in the 231:TKO cells expands the Myc-dependent transcriptome we identified in parental MDA-MB-231 cells. This expansion of Myc-dependent transcription following TXNIP loss occurs without an apparent increase in Myc’s intrinsic capacity to activate transcription and without increasing Myc levels. Together, our findings suggest that TXNIP loss mimics Myc overexpression, connecting Myc genomic binding and transcriptional programs to the metabolic signals that control TXNIP expression.

## Introduction

TXNIP is an α-arrestin protein with several anti-proliferative functions, predominant among these is an activity as a negative regulator of glucose uptake and as a suppressor of multiple pro-growth signaling pathways (1-5). Consistent with a role in restricting cell growth, TXNIP expression is suppressed by multiple cancer-associated pro-growth signaling pathways (6-9) and its expression is typically low in tumors compared to adjacent normal tissue. Further, TXNIP expression decreases with increasing tumor grade and low TXNIP expression is correlated with poor clinical outcomes in several cancers (10-15). Whether low TXNIP expression supports cell growth by increasing glucose utilization, activating pro-growth pathways or whether other mechanisms also contribute is currently unknown.

Our previous work demonstrated that low TXNIP expression correlates with poor clinical outcomes in TNBC, but not in other breast cancer subtypes (12). Further, this correlation is more pronounced in patients with elevated expression of the c-Myc transcription factor (Myc henceforth), suggesting functional interaction(s) between TXNIP and Myc. Their crosstalk likely occurs at least at two levels. First, Myc drives expression of many glucose-dependent biosynthetic pathways (16-22), suggesting that low TXNIP expression (high glucose uptake) in combination with high Myc expression (high glucose use) helps cells match glucose availability and utilization. Second, Myc represses TXNIP expression by displacing the MondoA transcriptional activator from a shared E-box element located just upstream of the TXNIP transcriptional start site (12). Together, these findings suggest a feed-forward mechanism where Myc’s repression of TXNIP increases glucose uptake to support Myc-driven and glucose-fueled biosynthetic pathways.

Myc is implicated in more than 50% of human malignancy with elevated levels of transcriptionally active Myc being critical for its oncogenic function. Elevated Myc levels arise from many mechanisms including, but not limited to, transcriptional and translational mechanisms and protein stability (23, 24). Myc drives transcription as a heterodimer with Max, with Myc levels being limiting for the formation of Myc:Max complexes (25). The model that has emerged over the last several years is that at physiological levels Myc:Max complexes bind to high affinity E-box sequences in the promoters and enhancers of genes involved in housekeeping pathways that support cell growth, such as ribosomal biogenesis (26). At oncogenic levels, Myc:Max complexes invade lower affinity sites in the promoters and enhancers of genes that are associated with processes critical to cellular transformation, such as signaling pathways. Thus, increasing Myc levels expands the Myc-dependent transcriptome rather than simply increasing the expression of Myc-dependent transcripts (27-30).

In this report, we provide evidence that TXNIP loss drives a global increase in Myc genomic binding and drives gene expression programs enriched for known Myc targets. Surprisingly, this expansion of the Myc-transcriptome was not accompanied by an increase in Myc protein expression, suggesting that TXNIP loss leads to an increase in Myc’s specific activity as a transcription factor.

### Results

### TXNIP Loss is transcriptionally similar to Myc Overexpression

To better understand how TXNIP contributes to the aggressive behavior of TNBC, we performed RNA-sequencing on RNA isolated from two separate clones of TXNIP null MDA-MB-231 cells (231:TKO) and MDA-MB-231 cells (parental 231) (Fig 1A, S1A Fig). Using cutoffs of reads >5 counts, and an adjusted p-value (pAdj) ≤0.05, we identified 1050 and 742 genes whose expression was up- and down-regulated, respectively in response to TXNIP deletion. INHBB, KISS1, FOXA2, and ZNF704, were among the most highly downregulated genes, whereas AKR1C3, MT-ATP8, and G0S2 were the most highly upregulated genes (S1B Fig).

**Fig 1.**
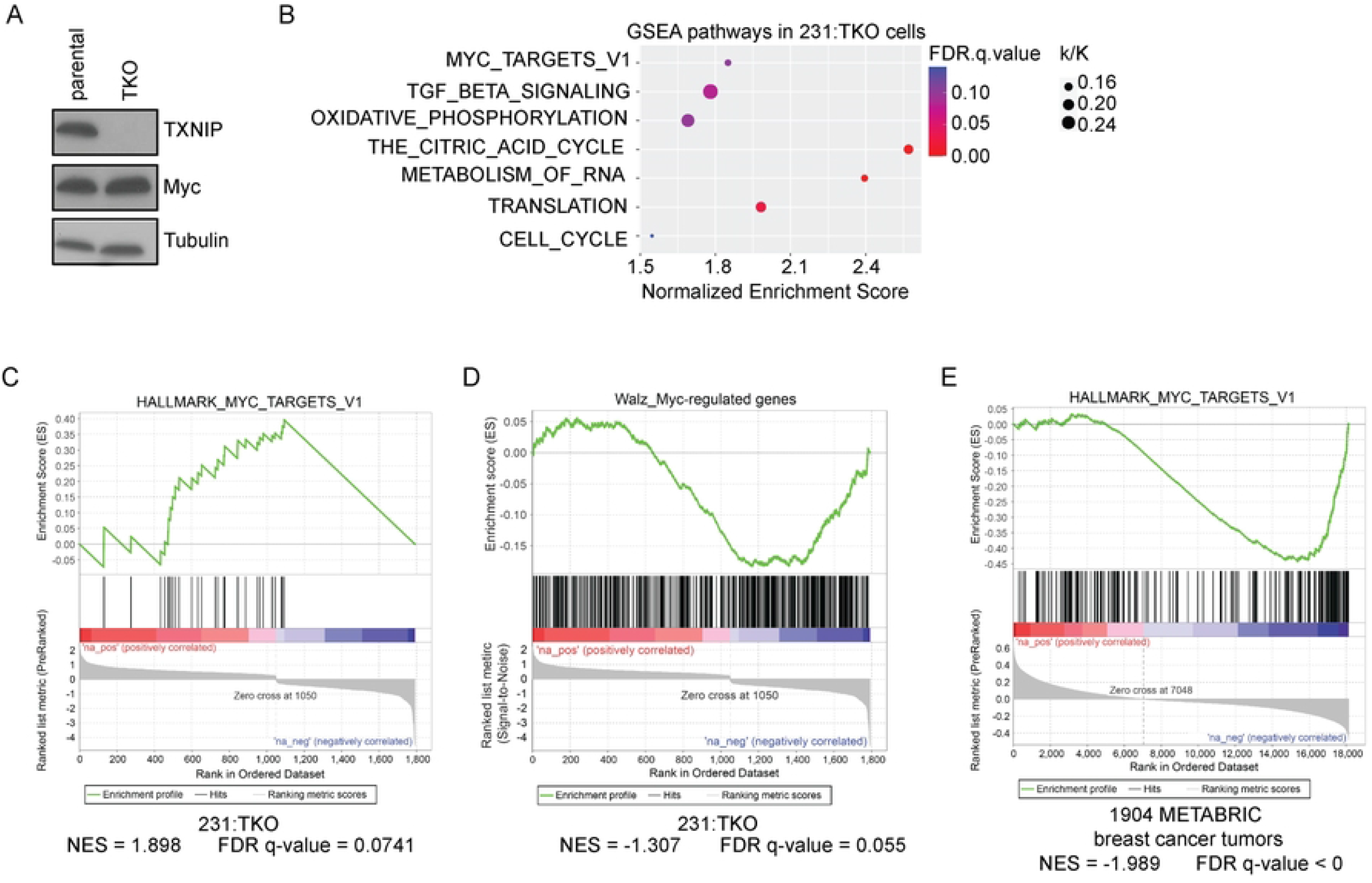
TXNIP loss is transcriptionally similar to Myc overexpression. (A) Western blotting was used to determine the levels of the indicated proteins in MDA-MB-231 (parental 231) and 231:TXNIP-knockout (TKO) cells. (B) RNA sequencing was performed on 2 biological replicates of each parental 231 and 231:TKO cells to identify TXNIP-regulated genes. Pre-ranked Gene Set Enrichment Analysis (GSEA) analysis was performed by comparing up- and down-regulated genes in 231:TKO cells with the Hallmark and Reactome datasets in the Molecular Signatures Database (MSigDB). The enriched GSEA pathways of TXNIP-regulated genes were plotted using ggplot2 package from R studio. k/K value is a ratio of number of genes in our data set (k) overlap with the number of genes in the indicated dataset (K). (C) The pre-ranked GSEA plot of enrichment of the regulated genes in 231:TKO cells with the HALLMARK_MYC_TARGETS_V1 dataset. (D) The pre-ranked GSEA analysis of our 231: TKO dataset with Myc-regulated genes (1.4-fold upregulated and downregulated genes) generated following inducible Myc expression in U2OS cells (GSE44672). (E) A rank ordered gene list was developed by correlating TXNIP expression with the expression levels of all transcripts expressed in the 1904 breast cancer tumors available in the Molecular Taxonomy of Breast Cancer International Consortium (METABRIC) dataset. This pre-ranked gene list was used in a GSEA analysis of the Hallmark datasets from the MSigDB. An enrichment plot for HALLMARK_MYC_TARGETS_V1 is shown.

To identify TXNIP-dependent pathways, we compared our 231: TKO dataset with annotated gene sets using pre-ranked Gene Set Enrichment Analysis (GSEA) (31, 32). The TXNIP-null dataset was positively enriched for known Myc targets and genes in pathways involved transforming growth factor beta (TGF-β) signaling, oxidative phosphorylation, the citric acid cycle, metabolism of RNA, translation, cell cycle, the G2M checkpoint and fatty acid metabolism (Fig 1B-C, S1C Fig). The identification of Myc targets and pathways known to be regulated by Myc in the 231:TKO cells raises the possibility that TXNIP loss resembles Myc overexpression. Importantly, the proliferation rate of 231:TKO cells is slightly slower than parental cells, suggesting that the in Myc transcriptional activity is not an epiphenomenon downstream of increased proliferation (S1D Fig).

Extending the correlation beyond breast cancer cell lines, we found that differentially regulated transcripts in 231:TKO cells were enriched for Myc-responsive transcripts in U2OS osteosarcoma cells (33) (Fig 1D). To experimentally validate our findings, we knocked TXNIP out in immortalized human myoblast MB135 cells (MB135:TKO) (34) (S1E Fig). We differentiated the parental MB135 and MB135:TKO cells into myotubes for 5 days and used RNA-seq to determine their transcriptional profiles. We found that differentiated MB135 cells were also enriched for known Myc targets (S1F Fig). Together these data suggest that TXNIP may be a general repressor of Myc transcriptional activity, capable of functioning in both cancer and immortalized cell lines from different lineages.

To determine whether the inverse relationship between TXNIP and Myc is restricted to cell lines, we generated a pre-ranked dataset by correlating *TXNIP* mRNA levels with the expression levels of all other transcripts expressed in the 1904 breast tumors annotated in the Molecular Taxonomy of Breast Cancer International Consortium (METABRIC) dataset (35, 36). This pre-ranked dataset was negatively enriched with a dataset containing known Myc targets (Fig 1E). This finding suggests that TXNIP-associated gene expression programs are negatively correlated with Myc-dependent transcriptional programs across human breast cancers. TXNIP-correlated gene expression programs were also negatively enriched with several other pro-growth datasets, including mTOR signaling, E2F targets, and additional Myc targets (S2A Fig), and positively enriched in several datasets including inflammatory responses, apoptosis, and adipogenesis (S2B Fig). Together, these data suggest that TXNIP may be a broad repressor of Myc transcriptional activity and function as a general negative regulator of cell growth.

**S1 Fig. TXNIP loss is transcriptionally similar to Myc overexpression**.

**Related to Fig 1**. (A) The relative *TXNIP* mRNA levels (normalized to that of β-actin) in parental 231 and 231:TKO cells were determined by RT-qPCR. (B) A volcano plot of the fold changes and adjusted p-values of regulated transcripts in 231:TKO cells. Gene expression changes in 231:TKO cells were determined using DESeq2. (C) A pre-ranked GSEA enrichment plots of the regulated transcripts in 231:TKO cells with the indicated Hallmark datasets. (D) Cell proliferation for parental 231 and 231:TKO cells in regular medium over a 94-hour time course was measured based on the percentage of confluency using real-time videography. (E) Western blotting was used to determine the levels of TXNIP protein and tubulin in parental MB135 and MB135:TKO cells. (F) A pre-ranked GSEA enrichment plot of regulated transcripts in differentiated myoblast MB135:TKO cells with the Hallmark_Myc_Targets_V1 dataset.

**S2 Fig. TXNIP-correlated gene expression programs are negatively correlated with pro-growth pathways**.

**Related to Fig 1**. (A) Transcripts positively correlated TXNIP expression across almost 2000 breast tumors are negatively correlated with genes in the 4 shown Hallmark datasets or (B) positively correlated with genes in the 4 shown Hallmark datasets.

### Gene expression changes in 231:TKO cells are Myc dependent

To determine whether the changes in gene expression identified in 231:TKO cells were Myc-dependent, we reduced Myc levels using a short interfering RNA approach (Fig 2A, S3A Fig). RNA-seq analysis revealed 5669 transcripts that were up- and down-regulated in 231:TKO+siMyc cells compared to 231+siNon-Targeting (siNT) cells using a pAdj<0.05. As expected, Myc was among the most down-regulated transcripts (S3B Fig) and the down-regulated genes in 231:TKO+siMyc cells were negatively enriched in known Myc targets (Fig 2B). The 231:TKO+siMyc dataset was also negatively enriched for known E2F targets and pathways involved in metabolism of RNA, protein translation, and cell cycle (Fig 2B, S3C Fig). Of the 1792 transcripts that were regulated in the 231:TKO cells, about 43% were (786 genes) were dependent on Myc (Fig 2C). Of these 786 genes, 548 transcripts (69.7%) were reciprocally regulated in 231:TKO and 231:TKO+siMyc cells, suggesting that TXNIP and Myc have primarily opposing functions in regulating gene expression (Fig 2D). Pathway analysis suggests that this group of reciprocally regulated genes largely account for the Myc-signatures identified in the 231:TKO+siMyc cells (Fig 2E). Ribosomal protein genes are well-established Myc targets (37), and they exemplify the reciprocal relationship between TXNIP and Myc. For example, TXNIP loss results in up-regulation of 24 ribosomal protein genes from both the large and small ribosomal subunit (30.4% of the 79 ribosomal protein genes) and all 24 of these genes were downregulated following Myc depletion (Fig 2F, S3D Fig). Collectively, these data show that TXNIP loss generates gene expression programs that not only resemble Myc overexpression, but that are also highly Myc-dependent.

**Fig 2.**
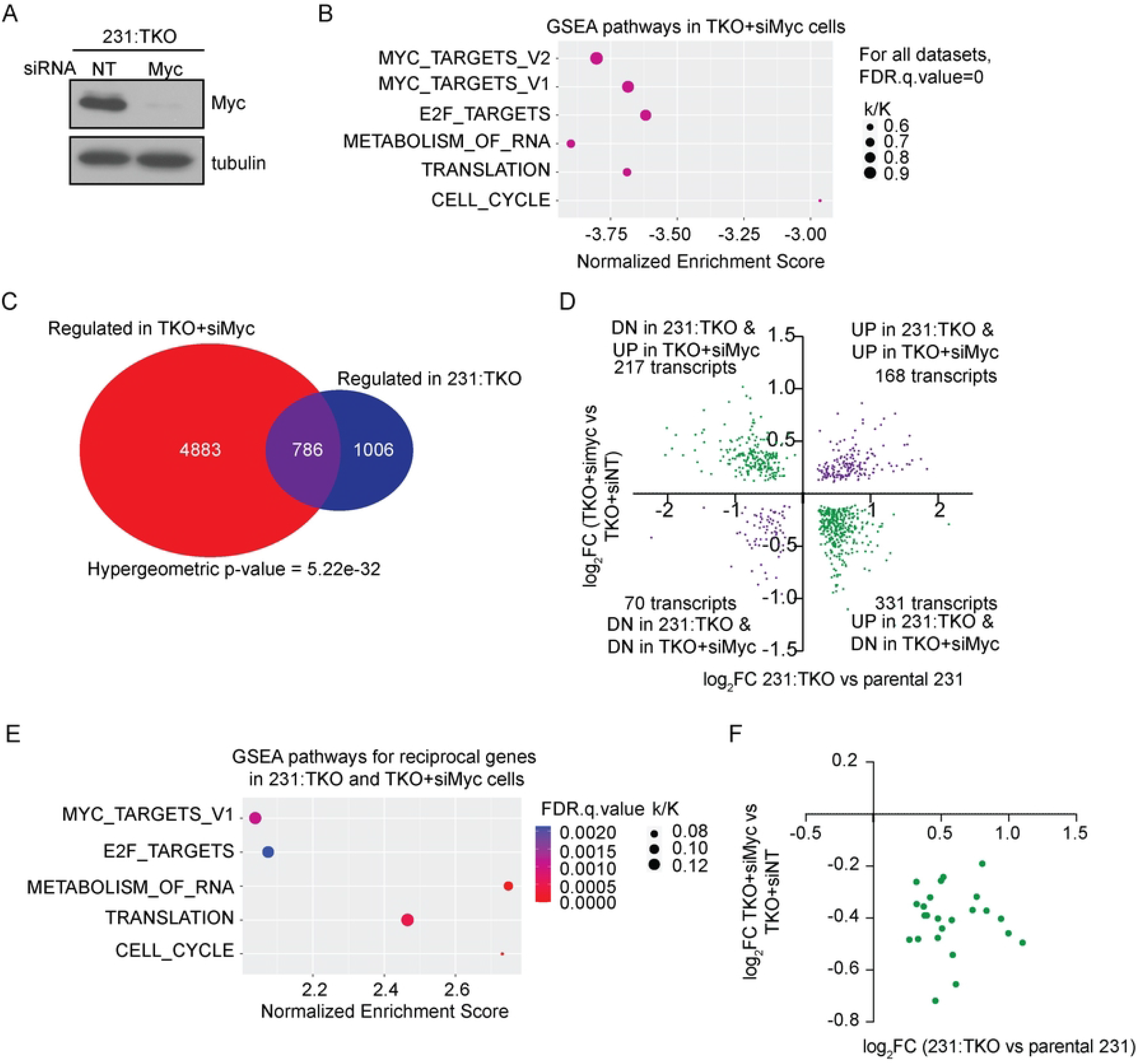
Gene expression changes in TKO are Myc dependent. (A) Western blotting was used to determine c-Myc protein and tubulin levels in 231:TKO cells following siRNA-mediated c-MYC knockdown. siRNA non-targeting control: siNT or siRNA targeting Myc: siMyc. (B) RNA sequencing was performed on 3 biological replicates 231:TKO with siNT or 231:TKO cells with siMyc to identify Myc-dependent genes in 231:TKO cells. Pre-ranked GSEA analysis was performed to identify pathways for Myc-dependent genes by comparing a ranked list of up- and down-regulated genes in 231:TKO+siMyc cells with the Hallmark and Reactome datasets in the MSigDB. The enriched GSEA pathways of c-Myc-dependent genes in 231:TKO cells were plotted using ggplot2 package from R studio. k/K value is a ratio of number of genes in our data set (k) overlap with the number of genes in the indicated datasets (K). (C) All regulated genes with adjusted p-value< 0.05 in 231:TKO were compared with all regulated genes with adjusted p-value <0.05 in 231:TKO+siMyc dataset to identify genes regulated in both datasets. The venn diagram was drawn using a VennDiagram package in R studio. (D) The 786 Myc-dependent transcripts genes were subdivided into 4 categories based the direction of their regulation in the 231:TKO and 231:TKO+siMyc datasets. (E) A ranked list of the 548 reciprocally regulated genes in (D) were used in a GSEA analysis using the MSigDB and the Hallmark and Reactome datasets. (F) Expression changes of 24 ribosomal protein transcripts regulated 231:TKO and 231:TKO+siMyc datasets.

**S3 Fig. Gene expression changes in 231:TKO cells are Myc dependent. Related to Fig 2**. (A) RT-qPCR was used to determine relative Myc mRNA levels (normalized to those of β-actin) in 231:TKO cells with siRNA non-targeting (siNT) or siRNA Myc-targeting (siMyc) treatment for 48 hours. (B) A volcano plot showing fold changes and adjusted p-values of transcripts differentially expressed in 231:TKO+siMyc cells. Differentially expressed genes were determined using DESeq2. (C) A pre-ranked GSEA was preformed using a ranked list of the differentially expressed genes in 231:TKO+siMyc cells and the Hallmark and Reactome datasets. (D) The log_2_ fold change in expression of six ribosomal protein genes in 231:TKO compared to their expression in parental 231 cells and in 231:TKO+siMyc cells compared to their expression 231:TKO+siNT cells. The gene expression changes all had an adjusted p-value (pAdj) of <0.05.

### TXNIP Regulates Global Myc Genomic Binding

Because TXNIP increased the expression of Myc transcriptional targets, we performed Myc ChIP-seq on parental 231 and 231:TKO cells to determine whether TXNIP regulates global Myc binding. We identified about 5600 Myc-occupied binding sites in parental 231 cells and roughly 28000 sites in 231:TKO cells (q-value cutoff =0.01) (Fig 3A), indicating that TXNIP is a broad repressor of Myc genomic binding. After filtering out counts with a percentage difference greater than 60%, we used the kmeans function in deepTools to identify 3 clusters of Myc genomic binding sites. In parental cells, Myc genomic binding was highest at sites in cluster 1 (669 sites), intermediate at sites in cluster 2 (3333 sites), and lowest at sites in cluster 3 (5063 sites). Myc binding in each cluster increased dramatically in 231:TKO cells. Genome browser views of sites in each cluster, RPL10A (cluster1), SLC18B1 (cluster 2), and NUP43 (cluster 3) showed a clear increase in Myc binding in 231:TKO cells as expected (Fig 3B). We also determined the fold increase or decrease in Myc binding (Myc signal ratio) in the 231:TKO cells for sites in each of the three clusters (Fig 3C). This analysis revealed that: 1) ∼96% of Myc binding sites showed elevated Myc occupancy in 231:TKO cells, 2) that the majority of Myc binding sites showed a slightly more than a 2-fold increase in Myc occupancy in the 231:TKO cells, and 3) that the increase in Myc signal ratio was similar for the binding sites in each cluster. Together, these data demonstrate that TXNIP loss leads to a global increase in Myc binding and support the hypothesis that TXNIP is a repressor of Myc genomic binding.

**Fig 3.**
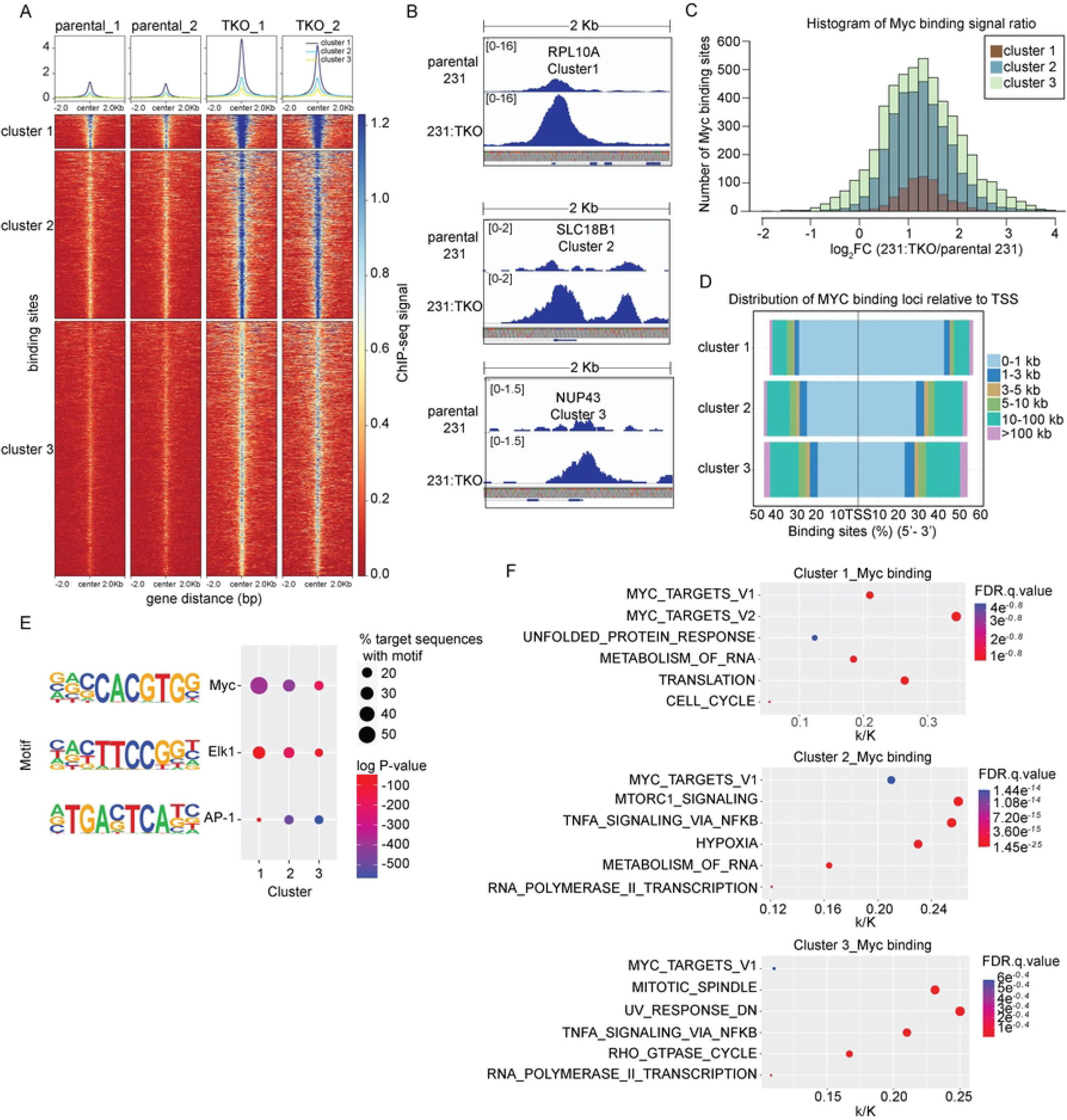
TXNIP regulates global Myc genomic binding. (A) A heatmap of Myc ChIP-sequencing data of 2 biological replicates of each parental 231 and 231:TKO cells was divided into 3 clusters using deepTools with the clustering argument of kmeans. (B) Myc binding, as visualized using IGV_2.5.2, on selected genes from each cluster in parental 231 and 231:TKO cells. (C) Histogram showing the Myc occupancy signal ratio of Myc binding sites in each cluster. Myc binding ratio was calculated by divided the counts in 231:TKO with the counts in parental 231 cells. (D) The distance of Myc binding sites from TSS in each cluster was annotated using the ChIPseeker program. (E) Enriched sequence motifs in the proximity of Myc-occupied sites in the 3 clusters were determined using Hypergeometric Optimization of Motif EnRichment (HOMER). (F) Myc-binding events were associated with potentially regulated genes using ChIPseeker. This set of Myc-associated genes were then evaluated for their enrichment in the Hallmark or Reactome datasets in MSigDB using GSEA. k/K value is a ratio of number of genes in our data set (k) overlap with the number of genes in the indicated dataset (K).

We used ChIPseeker (38) to associate Myc genomic binding events with specific genes and identified 644, 2680, and 3974 genes in cluster 1, 2, and 3, respectively. In cluster 1, 71.0% of the Myc binding sites were located within +/-1 kilobase (kb) of the transcriptional start site (TSS) of the associated genes. By contrast, the percentage of genes with TSS-proximal Myc binding sites progressively decreased in clusters 2 and 3, with a concomitant increase in Myc binding events at sites located between 10 and 100 kb from the from the TSS (Fig 3D).

We used Hypergeometric Optimization of Motif EnRichment (HOMER) (39) analysis to identify sequence elements enriched close to the Myc binding site in each of the three clusters. This analysis revealed that canonical CACGTG Myc-binding motifs were associated with roughly 50% of the Myc binding events in cluster 1, with the percentage of the canonical sites decreasing in clusters 2 and 3. We also discovered Elk1/ETS, and AP-1 motifs enriched close to the Myc-binding peak in all three clusters, with Elk1/Ets motifs decreasing from cluster 1 to 3 and AP1 motifs increasing (Fig 3E). Together these data suggest that cluster 1 contains the highest affinity canonical Myc binding sites that are located primarily in the promoters of the associated genes. Further, the data suggest that Clusters 2 and 3 contain lower affinity Myc binding sites that diverge from the canonical Myc binding element and are located more distal to the TSS, likely in regulatory enhancers.

Because, previous publications indicate that Myc can regulate different subgroups of targets based on the affinity of the Myc binding (27, 28), we used the MSigDB (31, 32) to identify the pathways enriched for Myc-binding sites in each cluster. This analysis revealed established Myc targets in each cluster with the highest percent enrichment in the Myc_Targets_V1 geneset in cluster1 and 2 with the k/K value of 0.21. The enrichment of this dataset decreased in cluster 3 with k/K value of 0.11. The Myc_Targets_V2 geneset was only enriched in cluster 1 with a k/K value of 0.34.

Genesets associated with translation and the metabolism of RNA, including genes encoding several ribosomal proteins, were enriched in clusters 1 and 2. mTORC1 and Hypoxia-associated genesets were strongly enriched in cluster 2, whereas TNFA_signaling via NFKB, mitotic_spindle and Rho_GTPase_cycle genesets were highly enriched in cluster 3 (Fig 3F). Together these data demonstrate different sets of target genes are enriched in each cluster and, consistent with previous reports (27, 28), suggest that Myc occupancy of different sets of target genes appears to be dictated by Myc’s differential binding to sites in the regulatory elements of these targets.

### G0S2 is reciprocally regulated by TXNIP and Myc

We next examined the Myc and TXNIP-dependent regulation of G0S2 as a representative of their reciprocal function in gene regulation. We chose to examine G0S2 for three reasons: 1) it was among the most highly upregulated genes in the 231:TKO cells (S1B and S4A Fig), 2) a previous publication showed that G0S2 was upregulated in the livers of TXNIP knockout mice (40) and 3) G0S2 in an inhibitor of triglyceride breakdown, which may contribute to the high levels of triglycerides observed in TXNIP knockout mice (41, 42) We first confirmed that G0S2 RNA and protein were upregulated in 231:TKO cells (Fig 4A and 4B). Our Myc ChIP-seq experiment revealed that Myc binding increased in the G0S2 promoter just upstream of the G0S2 transcriptional start site. This region encompassed a AGCGTGctcagCGCGTG sequence, which had been previously implicated in the glucose-dependent induction of G0S2 expression (43) (S4B Fig). The Myc binding site in the G0S2 promoter was in Cluster 2, suggesting that it is a medium affinity binding site. We confirmed that Myc binding increased at the G0S2 promoter following TXNIP loss using a ChIP-PCR approach (Fig 4C). As a negative control, Myc binding did not increase at genomic region on chromosome 10 that lacks a Myc binding site (Fig 4D). These data suggest that the increase in Myc binding observed at the G0S2 promoter, and presumably other promoters, reflects a bona fide increase in Myc binding rather than a general opening of chromatin in the 231:TKO cells that might support broad Myc binding.

**Fig 4.**
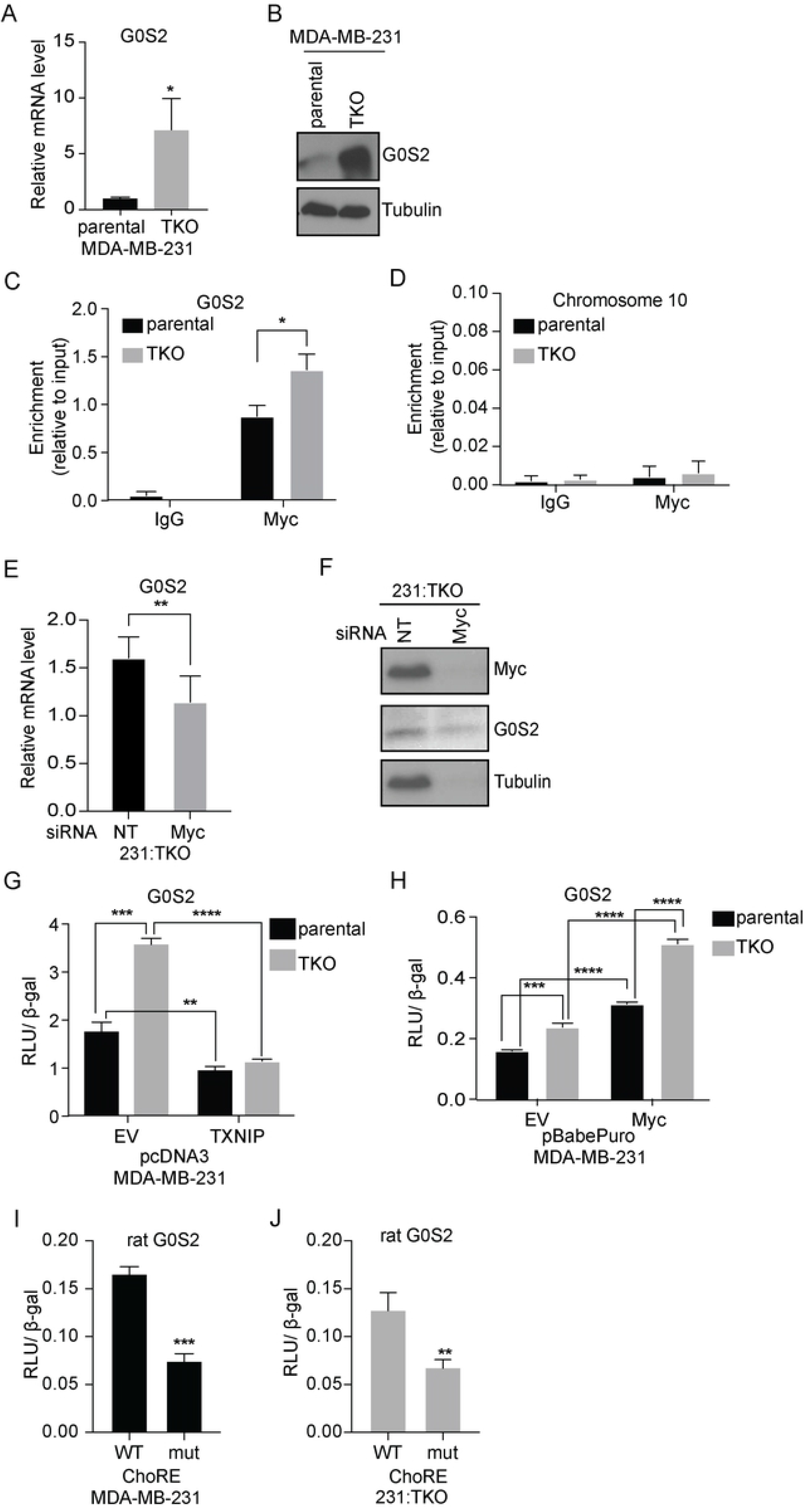
G0S2 is reciprocally regulated by TXNIP and Myc. (A) Human G0S2 (G0S2) mRNA levels in parental 231 and 231:TKO cells were measured using RT-qPCR. (B) Protein levels in parental 231 and 231:TKO cells were compared using Western blotting. (C and D) Three biological replicates of parental 231 and 231:TKO cells were used to perform Myc ChIP-qPCR to measure Myc occupancy upstream of the G0S2 transcriptional start site (C) and a region on chromosome 10 that lacks demonstrable Myc binding (D). Statistical significance was determined using a t-test. *p<0.05. (E and F) G0S2 mRNA (E) and protein (F) levels in 231:TKO cells were measured following siRNA-mediated c-MYC knockdown for 48 hours using RT-qPCR and Western blotting. (G) Luciferase activities of G0S2 luciferase reporter in lysates from parental 231 and 231:TKO cells with ectopic human TXNIP overexpression from pcDNA3 vector or pcDNA3 empty vector (EV) were measured. Luciferase activity was normalized to the beta-galactosidase (β-gal) activity. (H) Luciferase activities of G0S2 luciferase reporter in lysates from parental 231 and 231TKO cells with ectopic human Myc overexpression from pBabePuro vector or pBabePuro empty vector (EV) were measured. (I and J) The luciferase activities of wild-type (WT) rat G0S2-luciferase construct and mutated (mut) rat G0S2-luciferase construct in lysates from parental 231 (I) and 231:TKO (J) were measured. Carbohydrate response elements (ChoRE) of G0S2 promoter in the mutant rat G0S2-luciferase construct were deleted using site-directed mutagenesis (43). At least two biological replicates were carried out for all luciferase experiments. Representative figures were shown. Values are reported as mean and standard deviation. **p<0.01; ***p<0.001; ****p<0.0001.

The increase in Myc binding to the G0S2 promoter following TXNIP loss suggested that the elevated G0S2 expression in 231:TKO cells might be Myc-dependent. Consistent with this hypothesis, Myc knockdown in 231:TKO cells reduced the levels of G0S2 mRNA and protein (Fig 4E and 4F). To study the interplay between Myc and TXNIP in controlling G0S2 expression in more detail, we generated a luciferase reporter that contains 1493 base pairs of the human G0S2 promoter upstream of its translational start site. This fragment of the G0S2 promoter contains the Myc-occupied region identified in our ChIP-seq experiment. Consistent with TXNIP repressing G0S2 expression, the activity of the reporter was higher in 231:TKO cells than in 231 parental cells and overexpression of TXNIP suppressed reporter activity (Fig 4G). Further, Myc overexpression increased G0S2 reporter activity, suggesting that Myc is sufficient to activate G0S2 expression (Fig 4H).

Expression of rat G0S2 is glucose-dependent with glucose-responsiveness mapping to a Carbohydrate Response Element (ChoRE) just upstream of the TSS (43). Because ChoREs comprise two E-Box elements separated by 5 base pairs and the analogous region in the G0S2 promoter showed increased Myc occupancy in 231:TKO cells (S4B Fig), we tested whether TXNIP regulates G0S2 expression through the ChoRE. Like the human G0S2 promoter, the activity of the rat G0S2 promoter was elevated in 231:TKO cells and activity was blunted by overexpression of human TXNIP (S4C Fig). Deletion of the ChoRE in the rat G0S2 reporter reduced luciferase activities in the 231 parental and 231:TKO cells (Fig 4I and 4J), indicating that TXNIP regulated G0S2 expression through the ChoRE. Together these data validate the model that TXNIP loss leads to increased Myc binding and Myc-dependent activation of G0S2 gene expression.

**S4 Fig. G0S2 is reciprocally regulated by TXNIP and Myc**.

**Related to Fig 4**. (A) Genome browser view from RNA sequencing of human *G0S2* (G0S2) mRNA in parental 231 and 231:TKO cells. (B) Myc binding, as visualized using IGV_2.5.2, on G0S2 in parental 231 and 231:TKO cells. Putative carbohydrate response elements (ChoRE) in the G0S2 (43) was bound by Myc. (C) Luciferase activities of rat G0S2 reporter in parental and 231:TKO cells with ectopic human TXNIP overexpression in from pcDNA3 vector or pcDNA3 empty vector (EV) were measured. Luciferase activity was normalized to the beta-galactosidase (β-gal) activity. At least two biological replicates were carried out for all luciferase experiments. Representative figures were shown. Values are reported as mean and standard deviation. **p<0.01; ***p<0.001.

### TXNIP Controls Myc Transcriptional Programs by increasing Myc Binding

To examine the relationship between the Myc-dependent gene expression programs and increased Myc binding in 231:TKO cells, we compared the Myc-dependent transcripts identified in 231:TKO+siMyc cells with genes that showed increased Myc occupancy in 231:TKO cells. We found that 2903 (51.2%) Myc-dependent genes identified in 231:TKO+siMyc cells showed increased Myc binding in the 231:TKO cells (Fig 5A).

**Fig 5.**
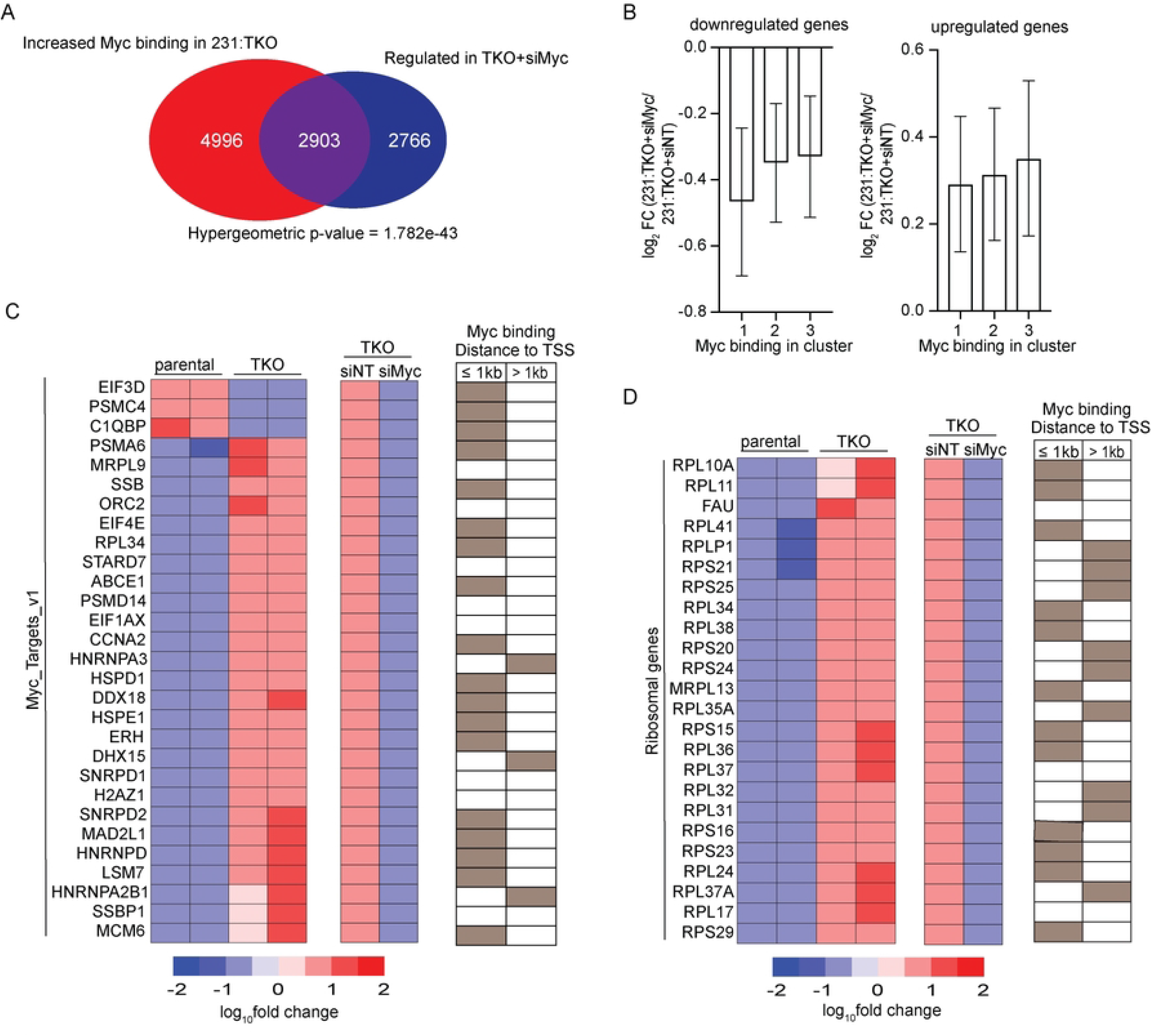
TXNIP controls Myc transcriptional program by increasing Myc binding. (A) Myc-dependent genes in 231:TKO cells were compared with genes that had increased Myc occupancy in 231:TKO cells. (B) The log_2_ fold change (log_2_ FC) of down-regulated and up-regulated genes in 231:TKO+siMyc cells compared to 231:TKO+siNT cells were plotted versus their Myc binding in the 3 Myc-binding clusters. (C and D) Heatmaps of regulated genes in 231:TKO cells and in 231:TKO+siMyc cells that are enriched in Myc_Targets_v1 (C) and in ribosomal protein genes (D) were plotted. Negative values of log_10_fold change indicate downregulation and positive values of log_10_fold change indicate upregulation. The distances of Myc binding sites from transcriptional start site (TSS) are determined using Genomic Regions Enrichment of Annotations Tool (GREAT) (44). Myc binding sites less than 1kb or more than 1kb from TSS are indicated by brown boxes. Open boxes indicate no Myc binding.

These 2903 genes were enriched in similar pathways as those enriched in 231:TKO+siMyc cells (S5A Fig). By contrast, we identified 4996 Myc binding sites that were not associated with changes in Myc-dependent gene expression in 231:TKO cells, suggesting that many Myc-binding events did not lead to measurable changes in gene expression. We examined two pathways of Myc-regulated genes in more detail. We found that 72.4% (21/29) and 87.5% (21/24) of the Myc-dependent transcripts enriched in the Myc_Targets_v1 gene set and genes encoding ribosomal proteins, respectively, showed elevated Myc binding close to the TSS (Fig 5C and 5D). Interestingly, most of the Myc_Targets_v1 had binding sites within 1kb of the TSS, whereas the proximity of the Myc binding site to the TSS of the ribosomal protein genes was more mixed with some genes having Myc binding sites within 1kb of the TSS, with others having sites more distant. In contrast to these genesets, transcripts in the oxidative phosphorylation pathway that were regulated by TXNIP loss, showed less Myc-dependence and fewer Myc-binding events (S5F Fig). These results suggest that TXNIP loss increases Myc-dependent gene expression by increasing Myc genome occupancy.

To better understand what constitutes a functional Myc binding event, we evaluated additional parameters. We found no correlation between the number of Myc sites and the magnitude of Myc-transcriptional regulation in either down- and up-regulated genes in 231:TKO+siMyc cells (S5B and S5C Fig). Further, the distance of the Myc-binding site relative to the TSS of a regulated gene did not correlate with the magnitude of Myc regulation. 60% of Myc-activated (i.e., downregulated in 231:TKO+siMyc) genes had a Myc binding site within 1kb of the TSS. By contrast, only 30% of the Myc-repressed genes (i.e., upregulated in 231:TKO+siMyc) had a Myc binding site within 1 kb of the TSS (S5D and S5E Fig). Although more Myc-activated genes had Myc binding sites closer to the TSS than Myc-repressed genes, there was no correlation between the degree of Myc regulation and the distance to the Myc binding sites for either Myc-activated or Myc-repressed genes loci. Finally, there was no relationship between the magnitude of Myc-dependence and whether the Myc binding site(s) associated with the regulated gene were present in cluster 1, 2, or 3 (Fig 3A and 5B). Thus, we observed that about 50% of the Myc-regulated genes had an associated Myc-binding event; however, there was no apparent relationship between the number of Myc binding sites, the affinity of those sites or the distance of the Myc binding site from the TSS and the magnitude of Myc-dependence.

**S5 Fig. TXNIP controls Myc transcriptional program by increasing Myc binding. Related to Fig 5**. (A) A list of 2903 genes that showed increased Myc binding in 231:TKO cells compared to parental 231 cells were ranked according to their differential expression in 231:TKO+siMyc cells. This list was analyzed using pre-ranked GSEA to identify enriched pathways in the MSigDB. (B and C) Differentially down-regulated (B) or up-regulated genes (C) in 231:TKO+siMyc cells were divided into groups based on the number Myc sites associated with each gene. (D and E) The distribution of Myc binding loci relative to the TSS for down-regulated (Myc-activated targets) (D) and up-regulated (Myc-repressed targets) (E) genes in 231:TKO+siMyc cells were annotated using ChIPseeker. The distance to the TSS was then compared change in gene expression following Myc knockdown. (F) Heatmap of genes regulated in 231:TKO cells that are enriched in the Reactome oxidative phosphorylation dataset. Differential regulation in 231:TKO+siMyc cells are indicated by yellow (upregulation) or green (downregulation) boxes. The distances of Myc binding sites from transcriptional start site (TSS) are determined using Genomic Regions Enrichment of Annotations Tool (GREAT) (44). The genes that have a Myc binding event within 1kb or more than 1kb from TSS are indicated in brown. Open boxes indicate no Myc binding.

### TXNIP Loss Expands the Myc Transcriptome

We next investigated whether TXNIP loss simply increased Myc-dependent transcriptional activity or whether its loss fundamentally altered the Myc-dependent transcriptome. We used an siRNA approach to knock Myc down in parental 231 cells and determined differentially expressed transcripts using RNA sequencing (S6A and S6B Fig). We identified 1196 genes that were Myc-dependent (pAdj < 0.05) in parental 231 and as expected, these genes were negatively enriched in Myc targets, E2F targets, and pathways involved in RNA metabolism, translation, and the cell cycle (Figure 6A).

**Fig 6.**
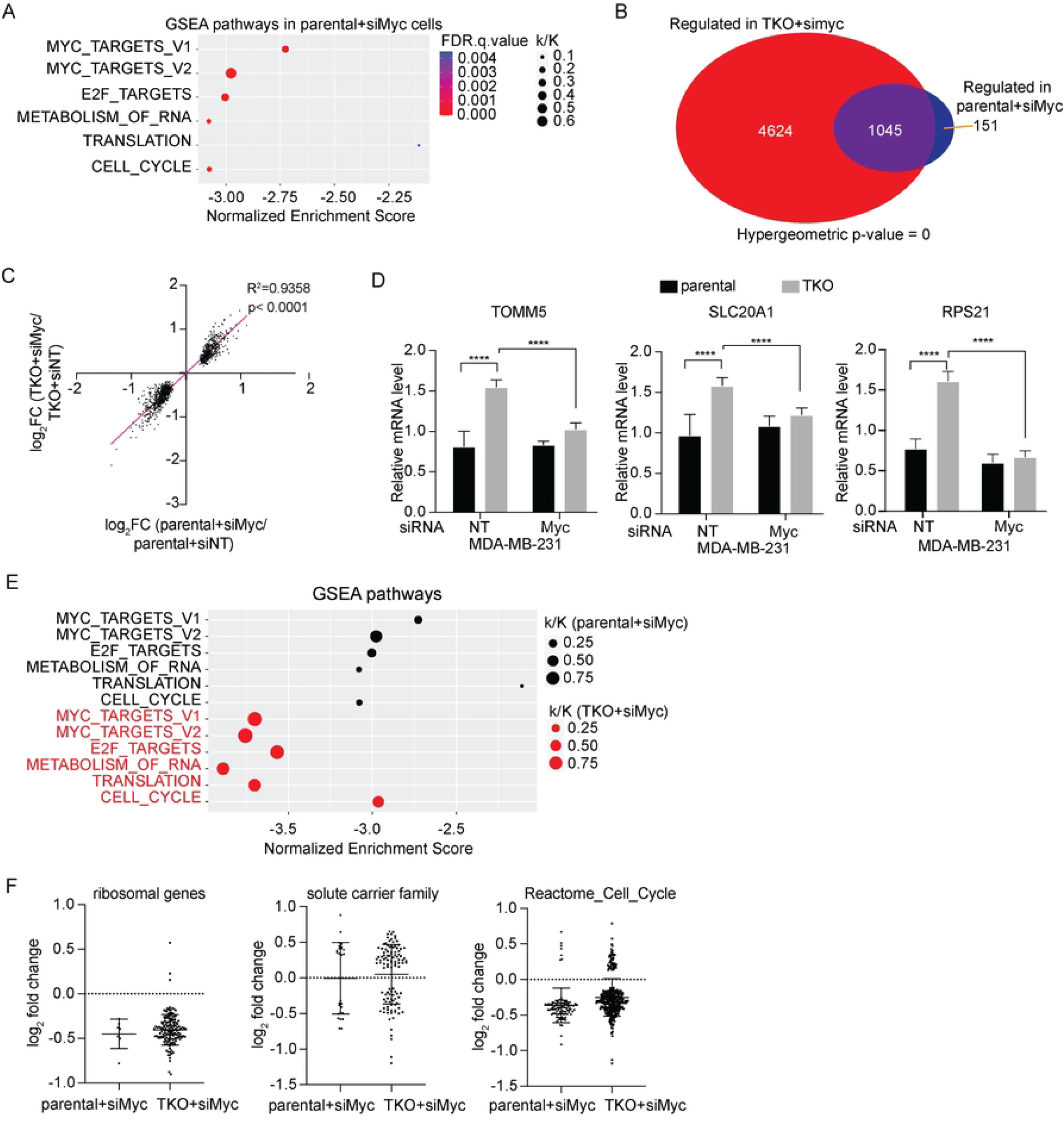
TXNIP loss expands the Myc transcriptome. (A) To identify Myc-dependent genes in MDA-MB-231 cells, RNA sequencing was performed on 3 biological replicates of each parental 231 with siRNA non-targeting control:siNT or with siRNA targeting Myc:siMyc. Pre-ranked GSEA analysis using a ranked list of Myc-dependent targets 231 parental cells and the Hallmark and Reactome datasets in the MSigDB. (B) Myc-regulated genes in parental 231 and in 231:TKO cells were compared. (C) The magnitude of the Myc-dependence of the 1045 transcripts that are Myc-dependent in both parental and 231:TKO cells was compared. (D) The Myc-dependence of 3 gene transcripts that were selected from the 4624 genes in Figure 6B were validated by reverse transcription-quantitative PCR (RT-qPCR). (E) Pre-ranked GSEA analysis was performed using ranked genes lists from Myc-dependent targets in parental 231 and 231:TKO cells and the Hallmark and Reactome data sets from the MSigBD. (F) The differential expression of genes in three different functional groups in 231 parental+siMyc and 231:TKO+siMyc datasets were compared.

The enrichment of these GSEA pathways was similar to that observed in 231:TKO+siyc cells, suggesting that, in general, Myc regulates similar gene expression programs in parental 231 and 231:TKO cells.

We next compared the Myc-dependent transcripts in parental 231 and 231:TKO cells. We first discovered that there were about 5 times as many Myc-dependent transcripts in 231:TKO cells (5669 transcripts) compared to the parental 231 cells (1196 transcripts) (Fig 6B), suggesting that TXNIP loss fundamentally alters the Myc-dependent transcriptome, rather than simply increasing Myc transcriptional activity. We next found that most Myc-dependent transcripts identified in the parental 231 cells (1045 transcripts/∼87%), also showed Myc-dependence in the 231:TKO cells. Interestingly, the magnitude of Myc dependence of these 1045 genes in the two cell populations was not significantly different (Fig 6C). There were 4624 Myc-dependent transcripts that were unique to the 231:TKO cells. We validated the expression of three transcripts, TOMM5, SLC20A1, and RPS21, that were part of this group. The levels of each transcript increased in 231:TKO cells and their expression was Myc dependent. By contrast, reducing Myc levels in parental 231 cells did not affect expression of any of these transcripts (Fig 6D). Together these findings suggest that TXNIP loss does not increase Myc transcriptional activity per se, rather TXNIP loss increases the number of Myc-dependent transcripts resulting in an expansion of the Myc-dependent transcriptome.

We conducted two additional analyses to validate the hypothesis that TXNIP loss expands the Myc-dependent transcriptome. First, we performed a pre-ranked GSEA analysis using the differentially expressed transcripts in the Myc-depleted parental 231 and 231:TKO cells. We found that each ranked transcript list was negatively enriched for known Myc targets, E2F targets and pathways involved in translation and the cell cycle. However, the Myc-regulated genes in 231:TKO cells showed lower normalized enrichment scores and increased overlap ratio (k/K) compared to parental 231 cells (Fig 6E), indicating that there were more Myc-regulated genes in these pathways in the 231:TKO cells. Finally, we used the Molecular Signature Database (MSigDB) (31, 32) to identify Myc-dependent transcripts associated with ribosomal function and the cell cycle and examined their Myc-dependence in parental 231 and 231:TKO cells. We also selected a group of transcripts encoding solute carrier proteins for evaluation. Even though there were many more Myc-dependent transcripts in each group in the 231:TKO cells, the degree of Myc-dependence is not significantly different between the two cell populations (Fig 6F). These results are consistent with the model that TXNIP loss broadens the Myc-dependent transcriptome to include additional transcripts associated with pathways that are well-documented to be Myc-regulated.

**S6 Fig. TXNIP loss expands the Myc transcriptome**.

**Related to Fig 6**. (A and B) RT-qPCR (A) and Western blotting (B) were used to determine Myc mRNA levels (normalized to that of β-actin) and levels of Myc protein in parental 231 with siRNA non-targeting (siNT) or siRNA Myc-targeting (siMyc) treatment for 48 hours.

## Discussion

We provide evidence that TXNIP is a broad and potent repressor of Myc genomic binding and transcriptional activity. TXNIP loss in an unrelated immortalized cell line also drove the gene expression programs containing known Myc targets and there is a strong negative enrichment for transcripts correlated with TXNIP expression across breast cancers and Myc-dependent gene signatures. These findings suggest that TXNIP’s ability to suppress Myc’s transcription function is not restricted to MDA-MB-231 cells, rather we speculate that TXNIP may be a general repressor of Myc-dependent transcription. Compared to the parental 231 cells, the Myc-dependent transcriptome is expanded in the 231:TKO cells (Fig 7A). This finding suggests that TXNIP loss does not simply increase the expression of the Myc-dependent transcripts present in the parental 231 cells but fundamentally alters Myc-driven transcriptional programs. Our ChIP-seq analysis suggests that the expanded Myc transcriptome in 231:TKO cells results from an increase in global Myc binding (Fig 7B). We show that the increase in Myc binding on G0S2 in 231:TKO cells results in upregulation of G0S2 expression. G0S2 is one example of target genes where TXNIP loss increases Myc-binding and transcriptional activity, but our analysis suggests that effect of TXNIP loss on Myc activity is global in nature and not restricted to G0S2. Our previous work showed that Myc can repress TXNIP expression by competing with its obligate transcriptional activator MondoA for a double E-box site in its promoter (12). Collectively, our findings here suggest that Myc-driven repression of TXNIP drives a feedforward regulatory circuit that reenforces Myc transcriptional programs.

**Fig 7.**
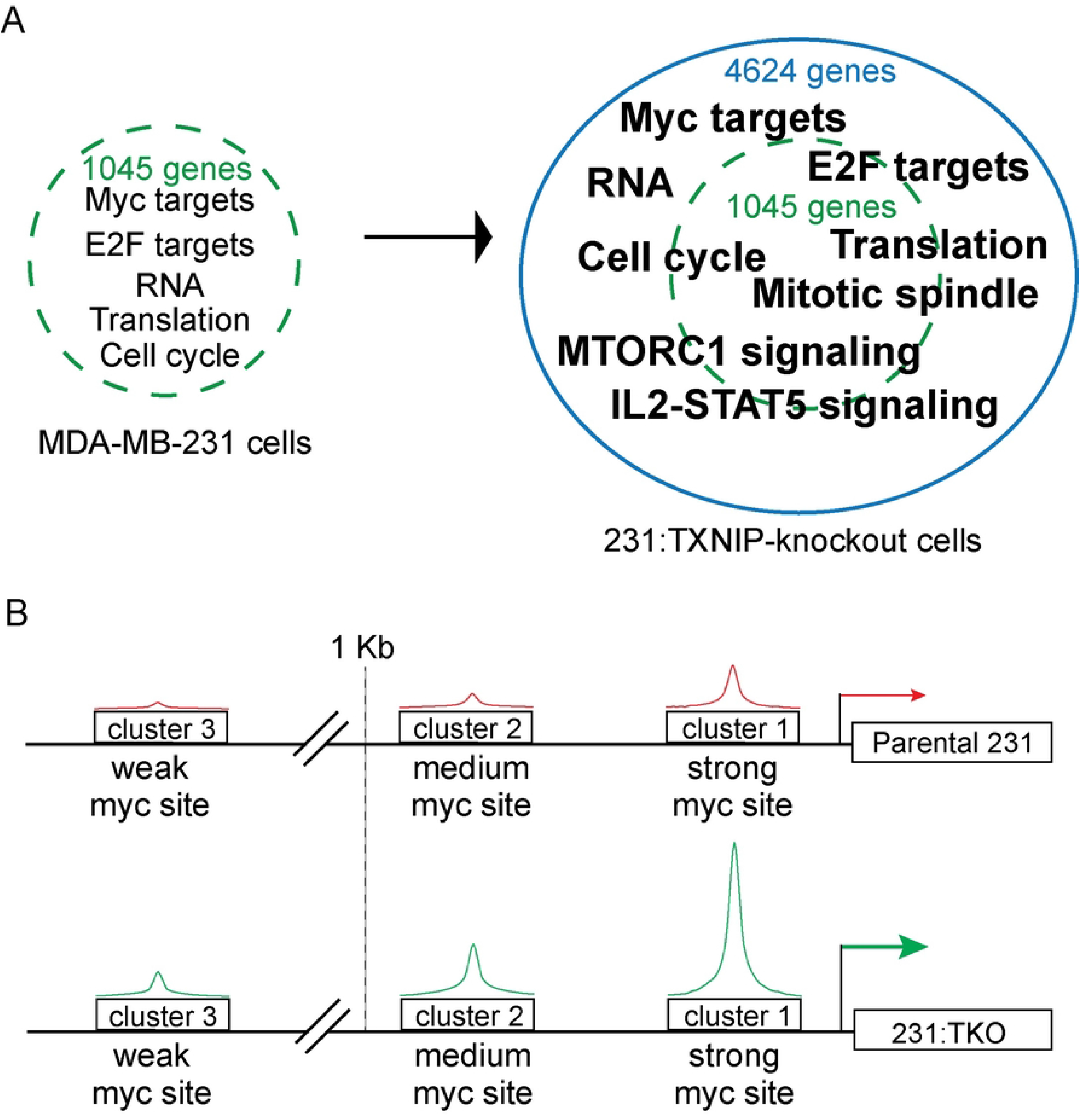
TXNIP is a repressor of Myc genomic binding and transcriptional activity. Following TXNIP loss the Myc-dependent transcriptome is expanded relative to that present in the parental MDA-MB-231 cells. Expansion in the 231:TKO cells does not dramatically alter the pathways regulated in the parental cells, rather there are more genes from a particular pathway regulated following loss of TXNIP. (B) We assigned Myc binding to three clusters based on whether they showed strong (cluster 1), medium (cluster 2) or weak (cluster 3) Myc binding in the parental 231 cells. The majority of sites in clusters 1 and 2 are located within 1kb of the transcriptional start site, whereas the majority of the sites in cluster 3 are more distal from the TSS and likely represent distal enhancers.

Control of TXNIP transcription, translation and stability is tightly coupled to progrowth signals. For example, mTOR blocks MondoA transcriptional activity, activated Ras blocks translation of TXNIP mRNA and AKT can phosphorylate TXNIP to trigger its degradation (7, 9, 45). One implication of these findings is that progrowth pathways via their suppression of TXNIP expression result in an indirect upregulation of Myc transcriptional activity. Another consequence of reducing TXNIP expression is an increase in glucose uptake, which we speculate provides carbon backbones for Myc-driven synthesis of macromolecules. This coordination of nutrient availability, i.e., low TXNIP levels and high glucose uptake, with nutrient use, i.e., Myc-driven synthesis of glucose-derived macromolecules, is likely important for supporting growth and proliferation of some cancer types. Supporting this hypothesis, a Myc_high_/TXNIP_low_ gene signature correlates with poor clinical outcomes in TNBCs, but not in other breast cancer subtypes (12).

Recent studies demonstrate that progressively increasing Myc levels drives Myc to low affinity non-canonical binding sites (27-30) and qualitative changes in Myc-dependent gene expression. The emerging model suggests that at low levels, Myc binds predominately to high affinity sites and regulates expression of genes that carry out essential housekeeping functions. As Myc levels increase, it invades lower affinity sites in promoter and enhancer regulatory regions. From these lower affinity sites, Myc is proposed to drive the expression of genes associated with its function as a transforming oncogene. The expanded Myc-dependent transcriptome in the 231:TKO cells mirrors these findings (Fig 7A-B). We divided Myc binding events in 231:TKO cells into high (cluster 1), medium (cluster 2), and low (cluster 3) affinity groups. The highest proportion of high affinity canonical CACGTG Myc binding sites were found in cluster 1, with the proportion decreasing in cluster 3 (Fig 3F). Myc binding events in cluster 3 were farther from the transcriptional start site, suggesting that TXNIP loss allows Myc to invade distal enhancer elements. Further, GSEA analysis revealed that high affinity Myc binding events in cluster 1 were associated gene sets enriched for housekeeping functions such as metabolism of RNA. By contrast, the sites in low affinity cluster 3 show enrichment for gene expression programs that may correspond to Myc’s function as a transforming oncogene, e.g., Rho signaling. The sites in medium affinity cluster 2 are enriched in gene sets associated with both Myc’s housekeeping and transformation-relevant transcriptional targets, suggesting an intermediate phenotype. With the increased level of Myc transcriptional activity in 231:TKO cells, particularly at potentially transformation-relevant targets, one might expect that they would display a higher level of cell growth-associated phenotypes. This does not to be the case at least in normal culture medium replete with glucose and serum-supplied growth factors (S1D Fig).

In general terms, Myc’s function in transcription and transformation is tightly linked to its absolute expression level (24). TXNIP loss expands Myc genomic binding and drives Myc-dependent gene expression programs, yet Myc protein levels do not increase following TXNIP loss (Fig 1A). Myc knockdown experiments showed that its intrinsic activity as a transcriptional activator is similar at Myc-dependent targets expressed in both parental 231 and 231:TKO cells (Fig 6C). In addition, the activity of multiple Myc-dependent luciferase reporters was similar in both cell types (unpublished data). By contrast, Myc genomic binding is dramatically increased in 231:TKO cells compared to parental cells with the increase in Myc binding similar in 231:TKO cells for for all three clusters. Together, these data suggest that TXNIP loss increases Myc’s intrinsic ability to bind genomic sites, rather than increasing its transcriptional activity per se. Although, we cannot formally rule out a direct role for TXNIP in regulating Myc genomic binding, we favor a model where TXNIP effects Myc genomic by an indirect mechanism. Our preliminary experiments provisionally rule out a role for TXNIP in regulating global chromatin accessibility, the amount of Myc in the nucleus, the formation of Myc:Max heterodimers, or Myc’s association with cofactors required for genome binding such as WDR5 (46); unpublished data). It is possible that TXNIP loss effects Myc genomic binding by altering its post-translational modification state or association with ancillary factors that stabilize its association with chromatin.

TXNIP deletion in parental 231 cells changes the expression of 1792 transcripts, yet the expression of only 786 transcripts is dependent upon Myc (Fig 2C). This suggests that in addition to regulating Myc genomic binding, TXNIP may regulate gene expression by additional mechanisms. TXNIP loss increases PI3K signaling, mTORC1 activity and alters cell metabolism (47-49), so there are several potential routes by which TXNIP loss may regulate gene expression independent of its effect on Myc activity characterized here. Alternatively, a recent report demonstrated that TXNIP loss can lead to a global up regulation in H3K27 acetylation (50), so an epigenetic mechanism is also possible.

This study and others demonstrate and establish that TXNIP is a repressor of at least two common features of the transformed state: Myc transcriptional activity and glucose uptake (1, 5, 51, 52) TXNIP expression is exquisitely dependent on the transcription factor MondoA (12, 53). Further, MondoA transcriptional activity seems to be primarily if not solely dedicated to regulating TXNIP (54). Together, these findings suggest that approaches to ectopically activate MondoA transcriptional activity might be useful cancer therapeutics in that they represent a way to block Myc transcriptional activity. For example, translation initiation inhibitors increase TXNIP expression in a MondoA-dependent manner (55). TXNIP induction is apparently independent of oncogenic burden, suggesting the potential utility of this approach.

## Acknowledgements

We thank members of the Ayer, Gertz and Varley labs for their thoughtful comments throughout this project and Elizabeth Leibold for comments on the manuscript. This work was funded by NIH grants, R01CA222650 (DEA) and R01 HG008974 (JG), and by 132596-RSG-18-197-01-DMC from the American Cancer Society (KEV). Funds from Huntsman Cancer Institute’s Cancer Center Support Grant (P30CA042014) and the Huntsman Cancer Foundation also provided support.

## Author Contributions

DEA, T-YL, JG, BRW and KEV designed the studies. T-YL, BRW, MLT, and KEM performed the experiments. DEA, T-YL, BRW, MEC and JMV performed data analysis. DEA and T-YL wrote the manuscript.

### Conflict of Interest

The authors declare no conflicts of interest

### Data availability

Raw and processed RNA-seq and ChIP-seq data has been deposited in the Gene Expression Omnibus under accession numbers GSE208412 and GSE208415

### Materials and Methods

#### Cell Culture Conditions

Parental MDA-MB-231, and 231:TKO cells were cultured in Dulbecco’s Modified Eagle Medium (DMEM) (Gibco; 1195073) with 10% Fetal Bovine Serum (FBS) (Gibco; A3160506), 1X MEM Non-Essential Amino Acids Solution (Gibco; 11140076) and 1X Penicillin-streptomycin (Gibco; 15140148). MB135 (34) and MB135:TKO cells were cultured in Ham’s F10 with L-glutamine (ThermoFisher; 11550043) with 20% FBS (Gibco; A3160506), 1X Penicillin-streptomycin (Gibco; 15140148), 10ng/ml recombinant human Fibroblast Growth Factor (Promega; G5071) and 1 μM dexamethasone (Sigma-Aldrich; D4902). For differentiation, MB135 and MB135:TKO cells were cultured in Ham’s F10 with L-glutamine (ThermoFisher; 11550043), 1X heat-inactivated horse serum (Sigma-Aldrich; H1270), 1X Penicillin-streptomycin (Gibco; 15140148), 10 μg/ml insulin from bovine pancreas (Sigma-Aldrich; I-1882) and 10 μg/ml transferrin (Sigma-Aldrich; T-0665). All cells were maintained at 37 °C and 5% CO2.

231:TKO cells were generated using human TXNIP CRISPR/Cas9 KO plasmid with TXNIP-specific guide RNA sequences from GeCKO (v2) library (Santa Cruz; sc-400664). TXNIP knockout clones were isolated from single cells and TXNIP knockout was validated using polymerase chain reaction (PCR) and western blotting using TXNIP (Abcam) antibodies. MB135:TKO cells were generated using human TXNIP CRISPR/Cas9 KO plasmid, three TXNIP-specific guide RNA sequences from GeCKO (v2) library (Santa Cruz; sc-400664) and a homology-directed repair (HDR) construct containing a puromycin-resistance cassette (Santa Cruz; sc-400664-HDR). TKO cells were isolated following selection of cells in 2.5 μg/mL puromycin. Loss of TXNIP was verified by immunoblotting.

#### Western Blotting

8×10^6^ cells were washed with 1X cold PBS once and scrapped with cell scrapper into ice-cold lysis buffer (400 mM NaCl, 20 mM HEPES [pH7.6], 1 mM EDTA, 1 mM EGTA, 25% glycerol and 0.1% NP-40) with protease inhibitors (1 mM PMSF, 2.5 μg/ml aprotinin, 1 μg/ml leupeptin and 1 μg/ml pepstatin), phosphatase inhibitor cocktail 1 (Sigma; P2850) and phosphatase inhibitor cocktail 2 (Sigma; P5726). Cells were disrupted using bioruptor (Diagenode; UCD-200) with a setting of 15 min, 30 seconds on, 30 seconds off. After sonication, disrupted cells were centrifuged at 14,000 rpm for 10 minutes. Supernatants were collected for further analysis. Protein concentrations were determined with a Bio-Rad protein assay (Bio-Rad; 5000006). Equivalent amounts of protein (40 – 80 μg) for different samples were resolved on SDS-PAGE, following transfer to PVDF membrane (Amersham; 10600023) with a setting of 150 V, 400 mA, and 1.5 hours at 4°C. After transfer, the PVDF membrane was blocked with 5% non-fat milk in 1X TBST (1X Tris-buffered saline, pH 7.4 with 0.1% Tween-20) for 1 hour.

Membranes were probed with primary antibodies using dilution between 1:500 and 1:2000 (TXNIP, Abcam, ab188865, 1:2000; c-MYC, Abcam, ab32072, 1:2000; G0S2, US Biological, 127066, 1:500 and alpha-tubulin, Molecular Probes, 236-10501, 1:20000) for overnight at 4°C. Protein signals were detected using HRP-conjugated mouse IgG (GE Healthcare, NA931, 1:5000), HRP-conjugated rabbit IgG (GE Healthcare, NA934, 1:15000) and ProSignal Pico ECL (Genesee Scientific, 20-300B).

#### Reverse Transcriptase Quantitative PCR (RT-qPCR)

RNA was extracted from cells using Zymo Research Quick RNA Miniprep Kit (Genesee Scientific, 11-328). 200 ng RNA was used to generate cDNA using GOScript Reverse transcriptase (Promega, A5001). qPCR was performed using CFX Connect Real-Time System and CFX Manager 3.1 program (Bio-Rad). Relative mRNA expression levels were determined from a standard curve generated for each RT primer set. Relative mRNA expression levels for different conditions/ samples were normalized to β-actin expression. Three biological replicates of experiments were performed. The values were reported as mean ± standard deviation of 3 technical replicates. Statistical significance was calculated using *t* test. Sequences of the primers used for RT-qPCR are listed in Table 1.

**Table 1.**
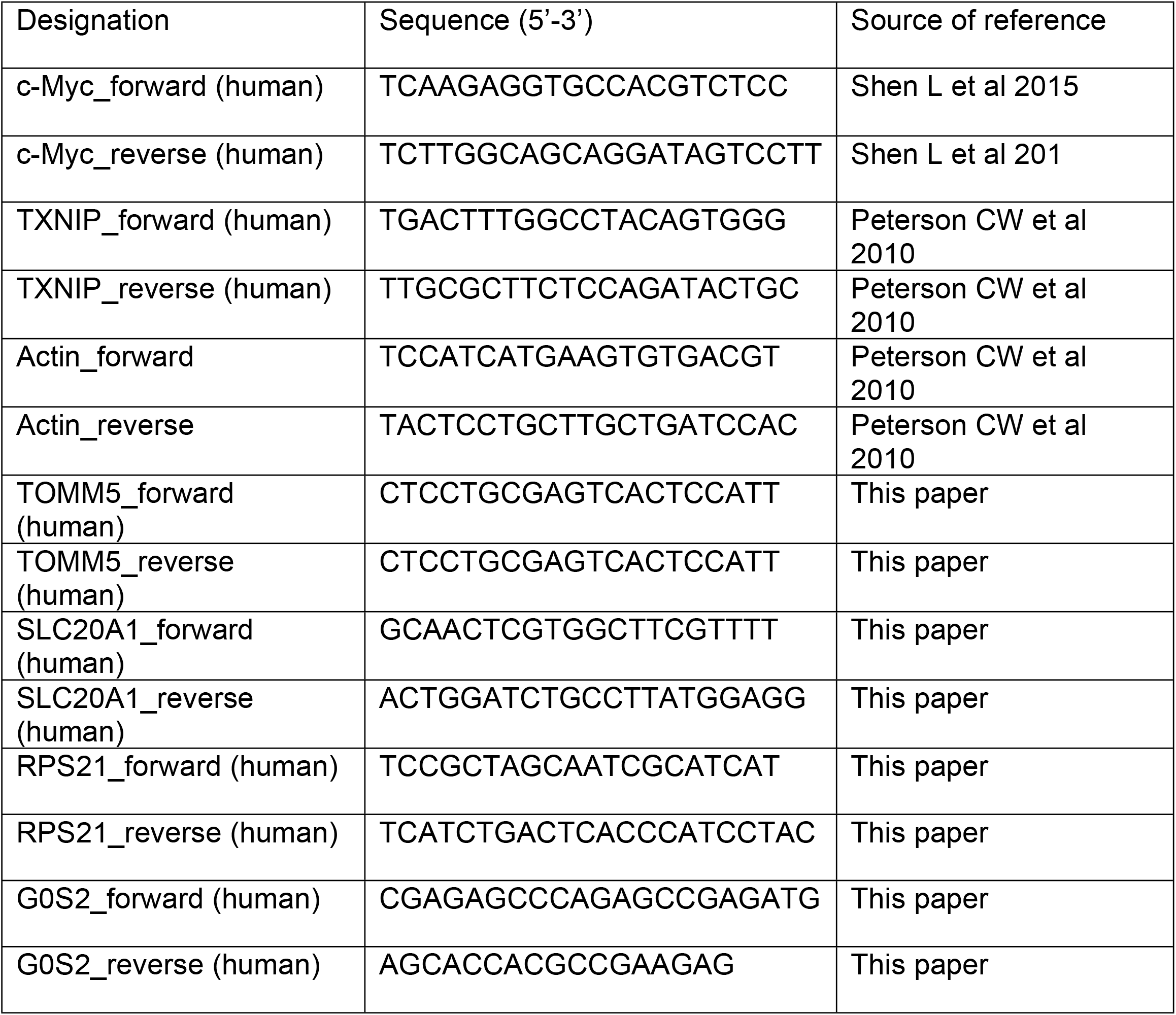
List of qRT-qPCR primers.

#### Chromatin Immunoprecipitation-Sequencing (ChIP-seq)

20×10^6^MDA-MB-231 or 231:TKO cells were cultured in DMEM (Gibco; 1195073) with 10% FBS (Gibco; A3160506), 1X MEM Non-Essential Amino Acids Solution (Gibco; 11140076) and 1X penicillin/streptomycin (Gibco; 15140148). Cells were crosslinked with 1% formaldehyde for 10 min at room temperature and then were treated with 0.125M glycine for 5 min to quench cross-linking reaction. Crosslinked cells were washed with 1X cold PBS and then were lysed in Farnham lysis buffer (5 mM PIPES pH 8.0, 85 mM KCl, 0.5% NP-40). Fixed chromatin was then harvested in Farnham lysis buffer (5 mM PIPES pH 8.0, 85 mM KCl, 0.5% NP-40) supplemented in protease and phosphatase inhibitors (ThermoFisher Scientific; A32959) by scraping off the plates with cells scrapers and transferred into 15 mL canonical tubes. Fixed chromatin was centrifuged at 2000 rpm for 5 min at 4°C and pellets were resuspended in RIPA buffer (1X PBS, 1% NP-40, 0.5% sodium deoxycholate, 0.1% SDS) for sonication. Sonication was performed on an Active Motif EpiShear Probe Sonicator for 5 min cycles of 30 sec on and 30 sec off with a 40% amplitude. Sonicated samples were centrifuged at 14000 rpm for 15 minutes at 4°C and supernatants were collected. Sonicated DNA, which had 200 to 500 base pair fragments was used for immunoprecipitation. For immunoprecipitation, 5 μg Myc (Cell Signaling; D3N8F) antibody was used. Anti-Myc was bound to Dynabeads M-280 sheep anti-rabbit (Invitrogen; 11204D) for at least 2 hours at 4°C. Bead-antibody complexes were incubated with fragmented chromatin overnight at 4°C with rocking. After overnight incubation, Dynabeads containing antibody-bound chromatin were captured with a magnetic rack. Dynabeads were washed with LiCl wash buffer (100 mM Tris [pH7.5], 500 mM LiCl, 1% NP-40 and 1% sodium deoxycholate) for 5 times at 4°C. Each wash was 3 minutes with rocking. After washing, the Dynabeads were washed once with 1 ml TE buffer (10 mM Tris-HCl [pH7.5] and 0.1 mM Na_2_EDTA). Beads were resuspended in 200 ml IP elution buffer (1% SDS and 0.1M NaHCO_3_) with vortexing. Beads were incubated in a 65°C bead bath for 1 hour with vortexing the tubes every 15 minutes to elute the antibody-bound chromatin from the beads. Beads were centrifuged at 14000 rpm for 3 minutes at room temperature. Supernatants (immune-bound chromatin) were collected and transferred to new microcentrifuge tubes. Input samples for each condition served as controls. Immuno-bound chromatin and inputs were de-crosslinked in a 65°C bead bath for overnight.

Reverse cross-linked DNA was cleaned up with ChIP DNA Clean & Concentrator Kit (Zymo Research; 11379C) according to the manufacturer’s protocol. Eluted DNA in EB buffer was used to construct ChIP library. Preparation of immunoprecipitated DNA for sequencing was performed as previously described (56). Briefly, blunted DNA fragments were ligated with sequencing adapters. The ligated DNA fragments were amplified with library PCR primers that contain barcodes (NEBNext ChIP-Seq Library Prep Reagent for Ilumina) for 15 cycles. Amplified DNA libraries from the anti-Myc ChIP were sequenced using Illumina HiSeq Sequencing with 50 cycles of single read. The resulting Fastq files were aligned to the human genome (hg19) using NovoAlign. Peaks were called using Model-Based Analysis of ChIP-seq-2 (MACS2) (57) using a p-value cut-off of 0.01 and the mfold parameter between 5 and 50. The Heatmap was generated using deepTools 2.0 (58). MYC-bound genes were annotated using the R package ChIPseeker (38). Myc binding motifs were determined using Hypergeometric Optimization of Motif EnRichment (HOMER) (39). GSEA and pre-ranked GSEA analyses (31, 32) were used to determine the pathway enrichment of Myc-bound genes. ggplot2 (59) was used to draw dot plots for pathway enrichment.

#### Chromatin Immunoprecipitation Quantitative PCR (ChIP-qPCR)

Anti-Myc immunoprecipitations were performed from chromatin isolated from 20×10^6^ MDA-MB-231 or 231:TKO cells as described above. Myc binding levels on genes were assessed with qPCR. qPCR was performed as described above. ChIP-qPCR primers that were used for experiments: human G0S2, forward:5’-TTCGCGTGCACACTGGCCTTCCC-3’, reverse: 5’-GAGGAGGGGAAAAGGAGGGGGTGGAAC-3’; human chromosome 10, forward:5’-GTCGGGAGCTTCCTATTCCTG-3’, reverse: 5’-AGAAGCCCACCCATCCCTAT-3’.

### RNA Sequencing Library Construction and Analysis

Total RNA was extracted from 3×10^6^ cells using the Zymo Research Quick RNA miniprep kit (Zymo Research; R1055). 500 ng of total RNA was used to capture mRNA and construct library using KAPA Stranded mRNA-Seq Kit (KAPA Biosystems; KK8420) according to the manufacturer’s protocol. Briefly, after mRNA capturing and fragmentation, cDNA was synthesized. Sequencing adaptors were ligated to the cDNA fragments. The adaptor-ligated cDNA fragments were amplified with library PCR primers that contain barcodes for about 10 cycles. Libraries were sequenced using either Illumina HiSeq 50 cycles Single-Read Sequencing or Novaseq Paired-Read Sequencing. The resulting Fastq files were aligned to the human genome (hg38) using STAR (60). Counts were generated using FeatureCounts version 1.63 (61) with the arguments “-T 24 -p -s 2 –largestOverlap “and using the Ensemble Transcriptome build 102 for GRCh38. DESeq2 (62) was used or determine the differential gene expression in different samples or treatments. Raw counts, rlog values, and normalized counts for each sample or treatment were generated with the DESeq2 program. Counts ≤ 5 were filtered and an adjusted p-value less than or equal to 0.05 was used in the DESeq2 program for determining differential gene expression. GSEA and pre-ranked GSEA analyses (31, 32) were used to determine the pathway enrichment of Myc-bound genes. ggplot2 (59) was used to draw dot plots for pathway enrichment.

#### Proliferation assay

Parental 231 and 231:TKO cells were plated in Dulbecco’s Modified Eagle Medium (DMEM) (Gibco; 1195073) with 10% Fetal Bovine Serum (FBS) (Gibco; A3160506), 1X MEM Non-Essential Amino Acids Solution (Gibco; 11140076) and 1X Penicillin-streptomycin (Gibco; 15140148) in 6-well plates with 6000 cells per well for each parental and 231:TKO cells. Proliferation was monitored for 7 days on the Incucyte Zoom Live Cell Imaging Platform (Sartorius) with 10X magnification and images were captured at 2-hour intervals. Confluency for parental 231 and 231:TKO cells was measured.

## Notes

### Competing Interest Statement

The authors have declared no competing interest.

## References

1. Hui TY, Sheth SS, Diffley JM, Potter DW, Lusis AJ, Attie AD, et al. Mice lacking thioredoxin-interacting protein provide evidence linking cellular redox state to appropriate response to nutritional signals. The Journal of biological chemistry. 2004;279(23):24387–93.

2. Minn AH, Hafele C, Shalev A. Thioredoxin-interacting protein is stimulated by glucose through a carbohydrate response element and induces beta-cell apoptosis. Endocrinology. 2005;146(5):2397–405.

3. O’Shea JM, Ayer DE. Coordination of nutrient availability and utilization by MAX- and MLX-centered transcription networks. Cold Spring Harbor perspectives in medicine. 2013;3(9):a014258.

4. Peterson CW, Ayer DE. An extended Myc network contributes to glucose homeostasis in cancer and diabetes. FBL. 2011;16(6):2206–23.

5. Stoltzman CA, Peterson CW, Breen KT, Muoio DM, Billin AN, Ayer DE. Glucose sensing by MondoA:Mlx complexes: a role for hexokinases and direct regulation of thioredoxin-interacting protein expression. Proceedings of the National Academy of Sciences of the United States of America. 2008;105(19):6912–7.

6. Elgort MG, O’Shea JM, Jiang Y, Ayer DE. Transcriptional and Translational Downregulation of Thioredoxin Interacting Protein Is Required for Metabolic Reprogramming during G(1). Genes Cancer. 2010;1(9):893–907.

7. Kaadige MR, Yang J, Wilde BR, Ayer DE. MondoA-Mlx transcriptional activity is limited by mTOR-MondoA interaction. Molecular and cellular biology. 2015;35(1):101–10.

8. Wu N, Zheng B, Shaywitz A, Dagon Y, Tower C, Bellinger G, et al. AMPK-dependent degradation of TXNIP upon energy stress leads to enhanced glucose uptake via GLUT1. Molecular cell. 2013;49(6):1167–75.

9. Ye Z, Ayer DE. Ras Suppresses TXNIP Expression by Restricting Ribosome Translocation. Mol Cell Biol. 2018;38(20).

10. Cadenas C, Franckenstein D, Schmidt M, Gehrmann M, Hermes M, Geppert B, et al. Role of thioredoxin reductase 1 and thioredoxin interacting protein in prognosis of breast cancer. Breast cancer research : BCR. 2010;12(3):R44.

11. Park JW, Lee SH, Woo GH, Kwon HJ, Kim DY. Downregulation of TXNIP leads to high proliferative activity and estrogen-dependent cell growth in breast cancer. Biochem Biophys Res Commun. 2018;498(3):566–72.

12. Shen L, O’Shea JM, Kaadige MR, Cunha S, Wilde BR, Cohen AL, et al. Metabolic reprogramming in triple-negative breast cancer through Myc suppression of TXNIP. Proceedings of the National Academy of Sciences of the United States of America. 2015;112(17):5425–30.

13. Tome ME, Johnson DB, Rimsza LM, Roberts RA, Grogan TM, Miller TP, et al. A redox signature score identifies diffuse large B-cell lymphoma patients with a poor prognosis. Blood. 2005;106(10):3594–601.

14. Zhou J, Chng WJ. Roles of thioredoxin binding protein (TXNIP) in oxidative stress, apoptosis and cancer. Mitochondrion. 2013;13(3):163–9.

15. Zhou J, Yu Q, Chng WJ. TXNIP (VDUP-1, TBP-2): a major redox regulator commonly suppressed in cancer by epigenetic mechanisms. Int J Biochem Cell Biol. 2011;43(12):1668–73.

16. Miller DM, Thomas SD, Islam A, Muench D, Sedoris K. c-Myc and cancer metabolism. Clin Cancer Res. 2012;18(20):5546–53.

17. Kim JW, Zeller KI, Wang Y, Jegga AG, Aronow BJ, O’Donnell KA, et al. Evaluation of myc E-box phylogenetic footprints in glycolytic genes by chromatin immunoprecipitation assays. Molecular and cellular biology. 2004;24(13):5923–36.

18. Shim H, Dolde C, Lewis BC, Wu CS, Dang G, Jungmann RA, et al. c-Myc transactivation of LDH-A: implications for tumor metabolism and growth. Proc Natl Acad Sci U S A. 1997;94(13):6658–63.

19. Gao P, Tchernyshyov I, Chang TC, Lee YS, Kita K, Ochi T, et al. c-Myc suppression of miR-23a/b enhances mitochondrial glutaminase expression and glutamine metabolism. Nature. 2009;458(7239):762–5.

20. Wise DR, DeBerardinis RJ, Mancuso A, Sayed N, Zhang XY, Pfeiffer HK, et al. Myc regulates a transcriptional program that stimulates mitochondrial glutaminolysis and leads to glutamine addiction. Proceedings of the National Academy of Sciences of the United States of America. 2008;105(48):18782–7.

21. Osthus RC, Shim H, Kim S, Li Q, Reddy R, Mukherjee M, et al. Deregulation of glucose transporter 1 and glycolytic gene expression by c-Myc. J Biol Chem. 2000;275(29):21797–800.

22. Dang CV. MYC, metabolism, cell growth, and tumorigenesis. Cold Spring Harb Perspect Med. 2013;3(8).

23. Meyer N, Penn LZ. Reflecting on 25 years with MYC. Nat Rev Cancer. 2008;8(12):976–90.

24. Conacci-Sorrell M, McFerrin L, Eisenman RN. An overview of MYC and its interactome. Cold Spring Harb Perspect Med. 2014;4(1):a014357.

25. Blackwood EM, Eisenman RN. Max: a helix-loop-helix zipper protein that forms a sequence-specific DNA-binding complex with Myc. Science. 1991;251(4998):1211–7.

26. Stine ZE, Walton ZE, Altman BJ, Hsieh AL, Dang CV. MYC, metabolism, and cancer. Cancer discovery. 2015;5(10):1024–39.

27. Lorenzin F, Benary U, Baluapuri A, Walz S, Jung LA, von Eyss B, et al. Different promoter affinities account for specificity in MYC-dependent gene regulation. Elife. 2016;5.

28. Altman BJ, Hsieh AL, Sengupta A, Krishnanaiah SY, Stine ZE, Walton ZE, et al. MYC Disrupts the Circadian Clock and Metabolism in Cancer Cells. Cell metabolism. 2015;22(6):1009–19.

29. Elkon R, Loayza-Puch F, Korkmaz G, Lopes R, van Breugel PC, Bleijerveld OB, et al. Myc coordinates transcription and translation to enhance transformation and suppress invasiveness. EMBO Rep. 2015;16(12):1723–36.

30. Sabò A, Kress TR, Pelizzola M, de Pretis S, Gorski MM, Tesi A, et al. Selective transcriptional regulation by Myc in cellular growth control and lymphomagenesis. Nature. 2014;511(7510):488–92.

31. Mootha VK, Lindgren CM, Eriksson KF, Subramanian A, Sihag S, Lehar J, et al. PGC-1alpha-responsive genes involved in oxidative phosphorylation are coordinately downregulated in human diabetes. Nat Genet. 2003;34(3):267–73.

32. Subramanian A, Tamayo P, Mootha VK, Mukherjee S, Ebert BL, Gillette MA, et al. Gene set enrichment analysis: a knowledge-based approach for interpreting genomewide expression profiles. Proc Natl Acad Sci U S A. 2005;102(43):15545–50.

33. Walz S, Lorenzin F, Morton J, Wiese KE, von Eyss B, Herold S, et al. Activation and repression by oncogenic MYC shape tumour-specific gene expression profiles. Nature. 2014;511(7510):483–7.

34. Stadler G, Chen JCJ, Wagner K, Robin JD, Shay JW, Emerson Jr CP, et al. Establishment of clonal myogenic cell lines from severely affected dystrophic muscles - CDK4 maintains the myogenic population. Skeletal Muscle. 2011;1(1):12.

35. Curtis C, Shah SP, Chin SF, Turashvili G, Rueda OM, Dunning MJ, et al. The genomic and transcriptomic architecture of 2,000 breast tumours reveals novel subgroups. Nature. 2012;486(7403):346–52.

36. Pereira B, Chin S-F, Rueda OM, Vollan H-KM, Provenzano E, Bardwell HA, et al. The somatic mutation profiles of 2,433 breast cancers refine their genomic and transcriptomic landscapes. Nature Communications. 2016;7(1):11479.

37. Schlosser I, Holzel M, Murnseer M, Burtscher H, Weidle UH, Eick D. A role for c-Myc in the regulation of ribosomal RNA processing. Nucleic Acids Res. 2003;31(21):6148–56.

38. Yu G, Wang LG, He QY. ChIPseeker: an R/Bioconductor package for ChIP peak annotation, comparison and visualization. Bioinformatics. 2015;31(14):2382–3.

39. Heinz S, Benner C, Spann N, Bertolino E, Lin YC, Laslo P, et al. Simple combinations of lineage-determining transcription factors prime cis-regulatory elements required for macrophage and B cell identities. Mol Cell. 2010;38(4):576–89.

40. Oka S, Yoshihara E, Bizen-Abe A, Liu W, Watanabe M, Yodoi J, et al. Thioredoxin binding protein-2/thioredoxin-interacting protein is a critical regulator of insulin secretion and peroxisome proliferator-activated receptor function. Endocrinology. 2009;150(3):1225–34.

41. Yang X, Lu X, Lombès M, Rha GB, Chi YI, Guerin TM, et al. The G(0)/G(1) switch gene 2 regulates adipose lipolysis through association with adipose triglyceride lipase. Cell Metab. 2010;11(3):194–205.

42. Bodnar JS, Chatterjee A, Castellani LW, Ross DA, Ohmen J, Cavalcoli J, et al. Positional cloning of the combined hyperlipidemia gene Hyplip1. Nat Genet. 2002;30(1):110–6.

43. Ma L, Robinson LN, Towle HC. ChREBP*Mlx is the principal mediator of glucose-induced gene expression in the liver. The Journal of biological chemistry. 2006;281(39):28721–30.

44. McLean CY, Bristor D, Hiller M, Clarke SL, Schaar BT, Lowe CB, et al. GREAT improves functional interpretation of cis-regulatory regions. Nat Biotechnol. 2010;28(5):495–501.

45. Waldhart AN, Dykstra H, Peck AS, Boguslawski EA, Madaj ZB, Wen J, et al. Phosphorylation of TXNIP by AKT Mediates Acute Influx of Glucose in Response to Insulin. Cell Rep. 2017;19(10):2005–13.

46. Thomas LR, Wang Q, Grieb BC, Phan J, Foshage AM, Sun Q, et al. Interaction with WDR5 promotes target gene recognition and tumorigenesis by MYC. Molecular cell. 2015;58(3):440–52.

47. Hui ST, Andres AM, Miller AK, Spann NJ, Potter DW, Post NM, et al. Txnip balances metabolic and growth signaling via PTEN disulfide reduction. Proceedings of the National Academy of Sciences of the United States of America. 2008;105(10):3921–6.

48. Spaeth-Cook D, Burch M, Belton R, Demoret B, Grosenbacher N, David J, et al. Loss of TXNIP enhances peritoneal metastasis and can be abrogated by dual TORC1/2 inhibition. Oncotarget. 2018;9(86):35676–86.

49. Meuillet EJ, Mahadevan D, Berggren M, Coon A, Powis G. Thioredoxin-1 binds to the C2 domain of PTEN inhibiting PTEN’s lipid phosphatase activity and membrane binding: a mechanism for the functional loss of PTEN’s tumor suppressor activity. Arch Biochem Biophys. 2004;429(2):123–33.

50. Bechard ME, Smalling R, Hayashi A, Zhong Y, Word AE, Campbell SL, et al. Pancreatic cancers suppress negative feedback of glucose transport to reprogram chromatin for metastasis. Nat Commun. 2020;11(1):4055.

51. Parikh H, Carlsson E, Chutkow WA, Johansson LE, Storgaard H, Poulsen P, et al. TXNIP regulates peripheral glucose metabolism in humans. PLoS Med. 2007;4(5):e158.

52. Chutkow WA, Patwari P, Yoshioka J, Lee RT. Thioredoxin-interacting protein (Txnip) is a critical regulator of hepatic glucose production. The Journal of biological chemistry. 2008;283(4):2397–406.

53. Peterson CW, Stoltzman CA, Sighinolfi MP, Han K-S, Ayer DE. Glucose controls nuclear accumulation, promoter binding, and transcriptional activity of the MondoA-Mlx heterodimer. Molecular and cellular biology. 2010;30(12):2887–95.

54. Wilde BR, Ye Z, Lim TY, Ayer DE. Cellular acidosis triggers human MondoA transcriptional activity by driving mitochondrial ATP production. Elife. 2019;8.

55. Wilde BR, Kaadige MR, Guillen KP, Butterfield A, Welm BE, Ayer DE. Protein synthesis inhibitors stimulate MondoA transcriptional activity by driving an accumulation of glucose 6-phosphate. Cancer & metabolism. 2020;8(1):1–13.

56. Reddy TE, Pauli F, Sprouse RO, Neff NF, Newberry KM, Garabedian MJ, et al. Genomic determination of the glucocorticoid response reveals unexpected mechanisms of gene regulation. Genome Res. 2009;19(12):2163–71.

57. Zhang Y, Liu T, Meyer CA, Eeckhoute J, Johnson DS, Bernstein BE, et al. Model-based analysis of ChIP-Seq (MACS). Genome Biol. 2008;9(9):R137.

58. Ramírez F, Ryan DP, Grüning B, Bhardwaj V, Kilpert F, Richter AS, et al. deepTools2: a next generation web server for deep-sequencing data analysis. Nucleic Acids Research. 2016;44(W1):W160–W5.

59. Wickham H. ggplot2 : Elegant Graphics for Data Analysis. Cham: Springer International Publishing : Imprint: Springer,; 2016.

60. Dobin A, Davis CA, Schlesinger F, Drenkow J, Zaleski C, Jha S, et al. STAR: ultrafast universal RNA-seq aligner. Bioinformatics. 2013;29(1):15–21.

61. Liao Y, Smyth GK, Shi W. featureCounts: an efficient general purpose program for assigning sequence reads to genomic features. Bioinformatics. 2014;30(7):923–30.

62. Love MI, Huber W, Anders S. Moderated estimation of fold change and dispersion for RNA-seq data with DESeq2. Genome Biol. 2014;15(12):550

